# Linkage disequilibrium between single nucleotide polymorphisms and hypermutable loci

**DOI:** 10.1101/020909

**Authors:** Sterling Sawaya, Matt Jones, Matt Keller

## Abstract

Some diseases are caused by genetic loci with a high rate of change, and heritability in complex traits is likely to be partially caused by variation at these loci. These hypermutable elements, such as tandem repeats, change at rates that are orders of magnitude higher than the rates at which most single nucleotides mutate. However, single nucleotide polymorphisms, or SNPs, are currently the primary focus of genetic studies of human disease. Here we quantify the degree to which SNPs are correlated with hypermutable loci by examining a range of mutation rates. We use established population genetics theory to relate mutation rates to recombination rates and compare the theoretical predictions to simulations. Both simulations and theory agree that, at the highest mutation rates, almost all correlation is lost between a hypermutable locus and surrounding SNPs. The theoretical predictions break down as the mutation rate increases, and consequently differ widely from the simulated results. The simulation results suggest that some correlation remains between SNPs and hypermutable loci when mutation rates are on the lower end of the mutation spectrum. Consequently, in some cases SNPs can tag variation caused by some hypermutable loci. We also examine the linkage between SNPs and other SNPs and uncover ways in which the linkage disequilibrium of rare SNPs differs from that of hypermutable loci.

## Introduction

Missing heritability and hypermutable loci Mutation can take many forms, and can occur at vastly different rates across the human genome [1]. Like recombination, mutation can disrupt linkage between two loci. Linkage permits one genetic variant to act as a proxy for other genetic variants. This allows for an estimation of genetic effects using only non-causal variants. In population genetics the relationship between linkage and mutation has received only limited attention, and most models assume all mutation rates are small and negligible (e.g. [2, 3]). The potential for hypermutable loci to cause diseases and influence quantitative traits is now becoming apparent [1], but the relationship between hypermutation and linkage is only beginning to be explored [4–6].

Hypermutable regions composed of tandem repeats are of particular interest because of the way in which they mutate. Tandem repeats expand and contract in repeat number at a rate that is orders of magnitude higher than the rate of single nucleotide point mutations [7–10]. These regions are able to mutate new alleles and then revert to their original form, all while maintaining their ability to expand and contract. Therefore, not only are many of these loci highly polymorphic, but their alleles can often be identical-by-state and not identical-by-descent. Furthermore, tandem repeats are the most common hypermutable loci in the human genome [1, 7], and are often found in regions of functional significance [11].

The rates of expansion and contraction at tandem repeats are known to depend on the length of the tandem repeats, the size of the repeated subunit and the sequence composition. The most mutable are tandem repeats composed of short subunits, called microsatellites (also known as short tandem repeats, or simple sequence repeats). These repeats can have mutation rates up to 10^−2^ [7], but most have rates between 10^−3^ and 10^−5^ [8–10]. The most hypermutable microsatellites tend to have a high A/T content and have a large number of repeated subunits. Because long microsatellites have a tendency to contract more often than they expand [12], microsatellites undergo a lifecycle in which they are “born” and “die” in the genome over evolutionary time [8,13].

Tandem repeats composed of subunits greater than nine base-pairs are called minisatellites. Unlike microsatellites, these tandem repeats are not known for their extreme mutability. Their mutation rates are not as well documented [14], but a method to estimate their relative mutation rates is available [15]. Minisatellites are thought to expand and contract in repeat number through recombination [16], in contrast to microsatellites which mutate primarily through polymerase slippage and subsequent mismatch repair [7,17].

Tandem repeat alleles are associated with a range of human diseases [14,18]. Of these diseases, perhaps the most well known are caused by expanded microsatellites: Fragile-X disease caused by an expanded CGG repeat [19], and Huntingon’s disease caused by an expanded CAG repeat [20]. Both of these repeats are found in promoters, functional regions near the start of a gene. Promoters have a relatively high density of tandem repeats, suggesting that these hypermutable sequences may play a role in regulating gene expression [11,21].

Although tandem repeats are potential sources of heritable disease, recent attention has focused on single nucleotide polymorphisms (SNPs) for genetic association studies due to technology that allows them to be inexpensively and rapidly genotyped genome-wide. Common SNP variants can be used to measure genome-wide relatedness, and this relatedness can explain a moderate portion of the heritability for complex traits [22]. However, many SNP studies have failed to uncover variants with significant associations [23]. Furthermore, even SNPs with the strongest associations can only explain a small fraction of heritable genetic variation [24].

This lack of significant GWAS hits has been referred to as “missing heritability” [23,24], and the heritability still not explained by modeling all genome-wide SNPs simultaneously has been termed the “still-missing heritability” [25,26]. Tandem repeats have been hypothesized to be partially responsible for missing heritability [18,27], and may also be partially responsible for some of the still-missing heritability. Due to their high mutability, tandem repeats can mutate away from linkage with surrounding SNPs, and therefore SNP association studies are not expected to pick up all of the heritability caused by hypermutable variants. Studies using large numbers of tandem repeat loci have shown that tandem repeat variants are usually very weakly linked with surrounding SNPs [4–6]. These studies highlight how SNP data can be uninformative about hypermutable loci, supporting the hypothesis that hypermutable loci are sources of missing heritability. [4].

However, not all tandem repeat variants are weakly tagged by SNPs. A recent genome wide association study of amyotrophic lateral sclerosis (ALS) in the Finnish population [28] uncovered a locus of interest that, through a following familial study, led to the discovery that a microsatellite tandem repeat is a prevalent cause of familial ALS [29]. In the C9orf72 gene, expansion of a CCGGGG repeat in the first intron results in a dominant allele that causes ALS and can also cause frontal-temporal dementia [29]. The expanded repeat allele is in strong linkage disequilibrium with surrounding SNPs [28,30,31]. Studies of the associated haplotype reveal that the expanded repeat likely arose only once [30,31] and then spread around the globe, possibly along with Viking conquests [32]. This discovery demonstrates that tandem repeat diseases can be uncovered from SNP association studies.

The 5HTTLPR gene provides another example of how SNPs can be associated with functional tandem repeat variants. Variation in a minisatellite within the 5HTTLPR promoter may be associated with a range of personality phenotypes and neurological diseases [33,34]. Two SNPs adjacent to the promoter repeat are in strong linkage disequilibrium with the repeat alleles that have been associated with disease (*r*^2^=0.72; [34]). Together, these studies raise the possibility that more tandem repeat alleles can be uncovered as sources of disease using SNP data.

Although hypermutable loci are potential causes of disease and modifiers of complex traits, there is limited theoretical work analyzing linkage between a hypermutable locus and surrounding SNPs. The seminal work by Ohta and Kimura, [2] set the groundwork for understanding how mutation and recombination rates combine to affect linkage disequilibrium (LD) between two biallelic polymorphisms. However, their approximation assumes very low mutation rates. When mutation rates are not low, their approximation breaks down. The goal of the current work is to examine analytical approximations of LD derived by Ohta and Kimura at higher mutation rates, and using simulations, examine the accuracy of their approximation. For our results to be directly comparable to the results of Ohta and Kimura our analyses are limited to biallelic hypermutable loci, and thus we do not directly model multi-allelic tandem repeat loci. We discuss this potential limitation in the Discussion.

## Materials and Methods

### Theory relating linkage disequilibrium with mutation rates

We examine the linkage disequilibrium between a hypermutable locus, *A/a*, and an adjacent SNP marker, *B/b*, defined by the following mutation dynamics:

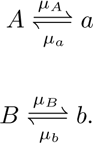

We model the hypermutable locus (A/a) as having only two alleles, with equal forward and backward mutation rates (so that *μ_A_* = *μ_a_*), although it does not perfectly correspond to hypermutable tandem repeat loci. This allows for a simple measure of correlation between the two loci, fitting the population genetics theory outlined below.

We assume the SNP locus (B/b) has a standard low mutation rate and the hypermutable locus has a high mutation rate, such that *μ_A_* + *μ_a_* >> *μ_B_* + *μ_b_*. The allele frequencies at locus *B* will be primarily influenced by drift, while the allele frequencies at *A* will be influenced by both drift and mutation (we ignore the possibility of selection). Denote the allele frequency of *A*(*B*) as *p_A_* (*p_B_*). The allele frequency at locus *A* is influenced by mutational equilibrium, in which:

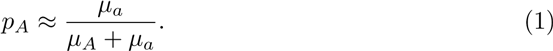

In a large population with limited drift, the frequency of allele *A* primarily depends on its forward and backward mutation rates. As population sizes get smaller, and/or the mutation rate gets lower, the allele frequencies are increasingly influenced by population dynamics (as shown in the results).

The allele frequencies at each locus are important because there is a relationship between the standardized measure of linkage disequilibrium (LD), *r*^2^, and relative allele frequencies [3, 35–38]. The maximum possible value of *r*^2^ between two loci is inversely related to the difference between the minor allele frequencies, so if there is a large difference in frequency between the two loci, *r*^2^ cannot be large [35,36,38].

Our primary interest is the expected correlation between two loci when one locus has a high mutation rate. For this, the frequency of haplotype AB will be defined as *p_AB_*. Linkage disequilibrium, *D*, is defined as:

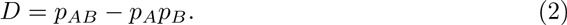

The square of the correlation between allele frequencies, *r*^2^, provides the proportion of variance at one locus that can be explained by another locus, and acts as a standardized measure of LD [3]:

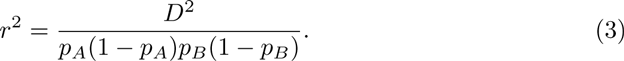

How much correlation is expected between loci? To examine this, [2] define a new variable, *ρ*^2^, as an approximation for *E*(*r*^2^). They use the approximation *E*(*x/y*) ≈ (*x*)/*E*(*y*) to find an approximation for *E*(*r*^2^),

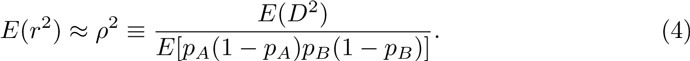

[2] then solve for the expected values of the numerator and denominator for a diffusion model, obtaining:

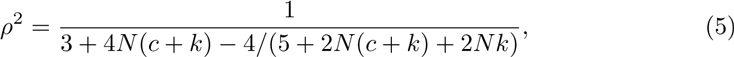

where N is effective population size, and *c* is the recombination rate between these two loci (here measured in centimorgans). The variable *k* is the sum of the mutation rates across both loci, *k* ≡ *μ_A_* + *μ_a_* + *μ_B_* + *μ_b_*, which is dominated by the mutation rates at the hypermutable locus (*k* ≈ *μ_A_* + *μ_a_*). To simplify notation, the forward/backward mutation rates at the hypermutable loci will be referred to as simply *μ*, such that *k* ≈ 2*μ*.

Somewhat counterintuitively, allele frequency is not present in the approximation for *ρ*^2^ (5). Although allele frequencies are present in the numerator, *E*(*D*^2^), and denominator, *E*[*p_A_*(1 − *p_A_*)*p_B_*(1 − *p_B_*)], their terms cancel resulting in an expression that only involves population size, N, recombination rate, *c* and the sum of mutation rates, *k* [2]. As discussed above, the maximum *r*^2^ value is determined by relative allele frequencies, but these results suggest that, on average at equilibrium, *r*^2^ is a function of only *N*, *c* and *k*. This prediction is examined here using simulated data (see next section). The simulations also use the diffusion model, so the equivalence of (4) and (5), as well as all of our results, rely on the assumptions of the model.

Furthermore, [2] showed that *ρ*^2^ is only an accurate approximation of *E*(*r*^2^) when *N*(*c* + *k*) is sufficiently larger than one. In this case *ρ*^2^ is approximated as:

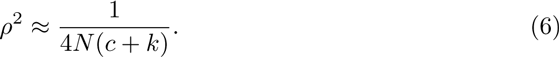

This approximation suggests that mutation and recombination act similarly to reduce linkage disequilibrium. Mutation is slightly different than recombination, however, because it changes allele frequencies, but this effect is reduced if the locus is in mutational equilibrium. More importantly, (6) also suggests that the expected correlation between allele frequencies is very small when *N*(*c* + *k*) is large. Therefore, if the mutation rate is large one would expect a weak correlation between a hypermutable locus and an adjacent SNP marker, unless the effective population size is small.

### 0.1 Simulations

Using the coalescent simulation program FastSimCoal [39], we simulated an effective population size of 10,000 individuals for a region of 100,000 base-pairs (100kb). At the center of the 100kb region we placed hypermutable locus (referred to as a “microsatellite” in FastSimCoal documentation) limited to only two alleles (A and a), with equal forward and backward mutation rates (*μ*) set to 10^−3^, 10^−4^, and 10^−5^ for different simulations. The use of FastSimCoal is relatively straightforward. To ease the reproduction of our simulations we provide the input simulation parameters and the random seed number in the supporting information file “S1 file”. Two thousand simulation results were obtained for each mutation rate. The recombination rate between adjacent base-pairs was set to 10^−8^, and the mutation rates at surrounding DNA loci were set to 5 · 10^−8^. The positions of the polymorphic locus, i.e. loci with a non-zero minor allele frequency, their variants, and the variants at the central hypermutable locus were retrieved from FastSimCoal. These results were converted to necessary file types using custom python scripts, and analyzed in python and R. There were 46 simulations for *μ* = 10^−5^ that were excluded because hypermutable loci were not polymorphic.

For each simulation, four statistics were calculated. First, the *r*^2^ values between the central hypermutable locus and surrounding SNPs were calculated. The mean of this value across simulations is referred to as “mean *r*^2^”. We expect this simulated measure of LD to be the most accurate estimate of the true degree of association because it does not rely on as many assumptions as the analytical approximation. Second, the average empirical values for *D*^2^ and *p_A_*(1 − *p_A_*)*p_B_*(1 − *p_B_*) were calculated from the simulations. We refer to the ratio of these two measures as “empirical *ρ*^2^”. Next, the values of *ρ*^2^ from (5) were calculated using the three parameters, *N*, *c*, and *k*, that were used in the simulation. We expect the analytical approximation *ρ*^2^ from (5) and empirical *ρ*^2^ to closely match because both the simulations and the statistical approach of [2] rely on the diffusion approximation. Finally, the position and *r*^2^ for the individual SNP with the highest *r*^2^ value were recorded from each individual simulation.

The simulation results were binned into regions of 100 base-pairs, corresponding to regions along the simulated chromosome relative to the position of the hypermutable locus. The values for *r*^2^ and empirical *ρ*^2^ were calculated and then averaged across SNPs for each 100 base-pair bin. The resulting plots were smoothed with LOESS smoothing.

To compare the hypermutable results with SNP-SNP correlations, we simulated a 150-kb region 50 times, with the same parameters as above (10,000 effective population 186 size, recombination rates of 10^−8^, and mutation rate of 5 · 10^−8^). For each simulation, we used SNPs that were at least 50-kb from the end of the region. Each SNP in this central region was examined separately for its correlations with surrounding SNPs at most 50kb away. This is equivalent to a central SNP in a 100kb region, thus making the LD between two SNPs comparable to the LD between SNPs and hypermutable loci.

## 1 Results

### 1.1 Allele frequencies from simulations

Fig. 1 (a)-(c) display the minor allele frequencies (MAFs) for the hypermutable loci, for each mutation rate. At mutation rates of 10^−3^ or 10^−4^ most of the hypermutable alleles have a high MAF. These high mutation rates drive the allele frequencies toward their mutational equilibria of 0.5. In contrast, the allele frequencies for loci with the mutation rate of 10^−5^ are strongly right skewed, with mostly rare alleles. At this lower mutation rate, the allele frequencies appear to be strongly influenced by population dynamics.

**Figure 1.**
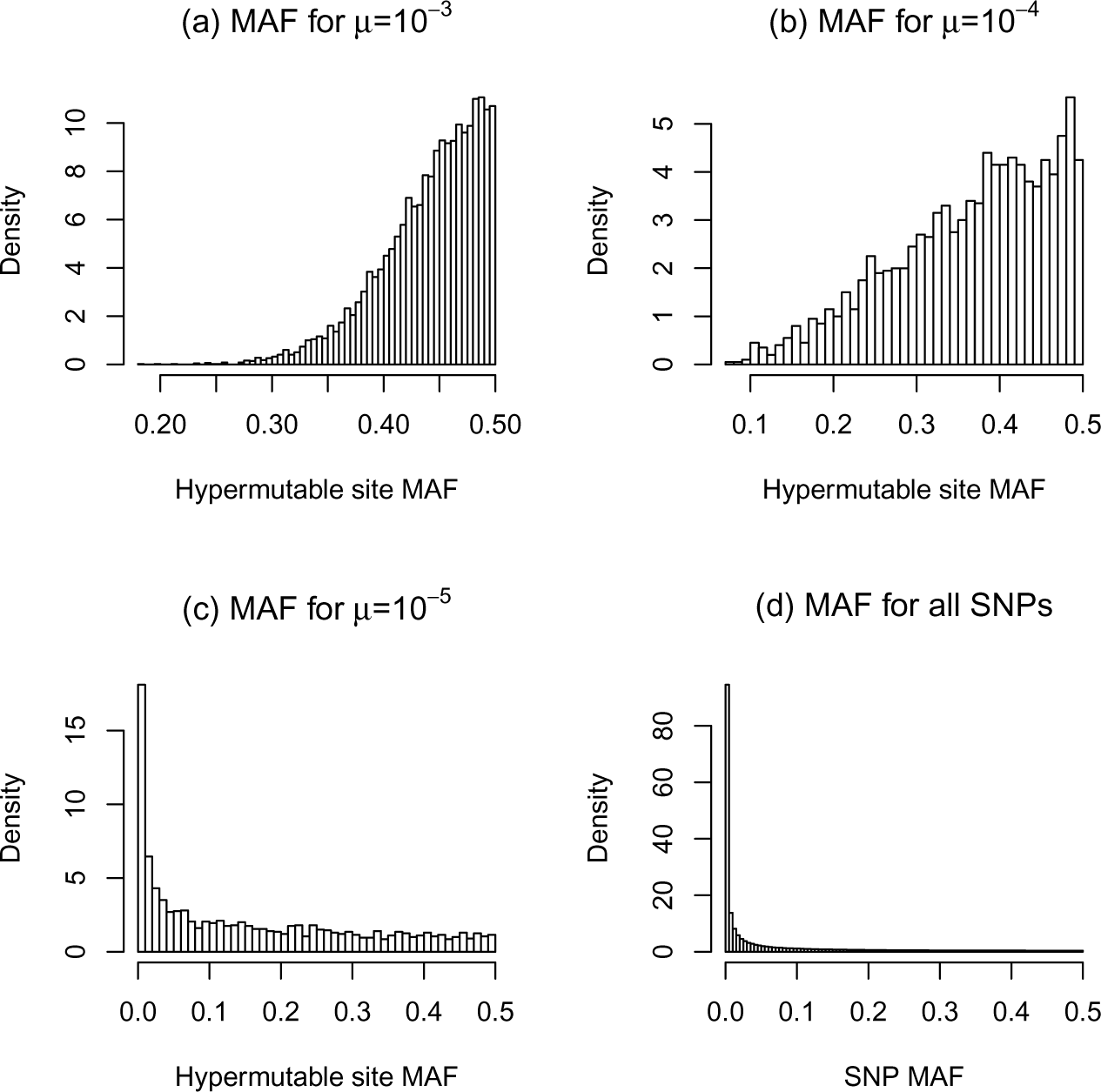
Histograms of allele frequencies from the simulations. The minor allele frequencies for bi-allelic hypermutable sites with mutation rates of 10^−3^ (a) 10^−4^ (b) and 10^−5^ (c) are shown. Only simulations with non-zero allele frequencies were used. Plot (d) shows a histogram of minor allele frequencies for SNPs in the simulation.

The simulated SNP allele frequencies are also strongly influenced by population dynamics, and the MAFs for most of these loci are very low (Fig. 1(d)). As discussed previously, the difference in allele frequencies between two loci influences their maximum possible *r*^2^. Hypermutable loci with a mutation rate of 10^−3^ have, on average, a high MAF, whereas the average SNP MAF is very low. Therefore, a large difference in allele frequencies exists between rare SNPs and most hypermutable loci, limiting their maximum *r*^2^.

### 1.2 Comparing *r*^2^ estimates with simulated results

For each mutation rate we plot the mean *r*^2^ between a central hypermutable locus and SNPs with any MAF across the entire simulated region (Fig. 2, green line). These mean *r*^2^ values are primarily influenced by associations between hypermutable loci and rare SNPs. The mean *r*^2^ values for simulations with a mutation rate of 10^−3^ are very low (Fig. 2 (c)), increasing slightly for 10^−4^ (Fig. 2 (b)), and more so for 10^−5^ (Fig. 2 (a)). We also plot the estimate of *ρ*^2^ made by [2], equation (5), in red. This approximation is greater than the mean *r*^2^ value for each scenario examined here, and much greater when the mutation rate is low or the inter-locus distance is short. Importantly, when mutation rates are low or loci are in close proximity, the value of *N* (*c* + *k*) is much less than 1. Consequently, as predicted by [2], this causes the estimate of *ρ*^2^ to differ from the mean *r*^2^.

**Figure 2.**
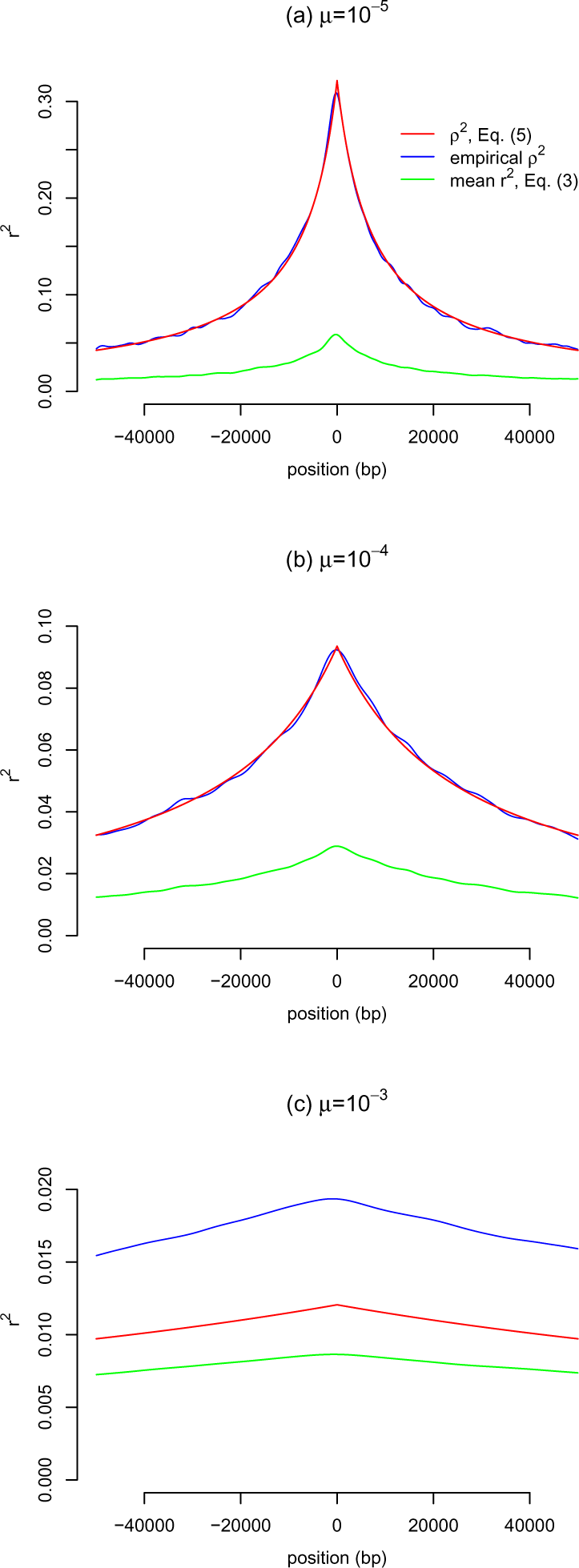
Plots comparing mean *r*^2^ from simulations (green), its approximation, *ρ*^2^ (red), and the empirical value of *ρ*^2^ (blue). The hypermutable locus is central (position 0), and *r*^2^ values were calculated between the central hypermutable element and surrounding SNPs. Results for simulations using hypermutable mutation rates of 10^−3^ (a), 10^−4^ (b), and 10^−5^ (c) are shown. The values of *ρ*^2^ are far greater than the mean *r*^2^, with the greatest difference found for low mutation rates. The values were calculated for bins of 100 base-pairs, and a line was drawn between these binned values using LOESS smoothing. Note the change in scale on the vertical axes between plots of different mutation rates.

Because the simulations use the same diffusion approximation assumptions as the analytical approach of [2], we expect the empirical *ρ*^2^ to match the approximation *ρ*^2^ from (5). Empirical *ρ*^2^ and the approximation (5) are nearly identical for the simulations using a hypermutable mutation rate of 10^−5^ or 10^−4^, but not for 10^−3^ (Fig. 2, blue and red lines). To examine whether the unexpected results for 10^−3^ were caused by a requirement for more simulations to converge to the analytical estimate, we ran an additional 8,000 simulations using this mutation rate. The results from all 10,000 simulations did not differ from the results of only 2,000 simulations (results not shown). We were therefore not able to determine the cause of this discrepancy, but nevertheless, for a mutation rate of 10^−3^ all three measures of *r*^2^ are very small.

Importantly, the mean *r*^2^ measured here uses hypermutable loci and SNPs with any allele frequencies above 0 (following the assumptions of [2]). This corresponds to a study in which all, or most, SNPs are genotyped, such as a sequencing study. If a study only uses common alleles, such as on a SNP chip with only common SNPs (MAF > 0.05), then the mean *r*^2^ values found between these common SNPs and a hypermutable site should be different.

To address how SNP minor allele frequencies influence the *r*^2^ between the SNPs and hypermutable loci, we examine the *r*^2^ values for SNPs with different MAFs, averaged across all regions. The horizontal black line in Fig. 3 shows the mean empirical *r*^2^ for SNPs binned by MAF value, for each mutation rate. The outer ends of the red vertical lines in this figure indicate the range between the 25th and 75th percentiles (5th and 95th for the ends of the thinner blue lines).

**Figure 3.**
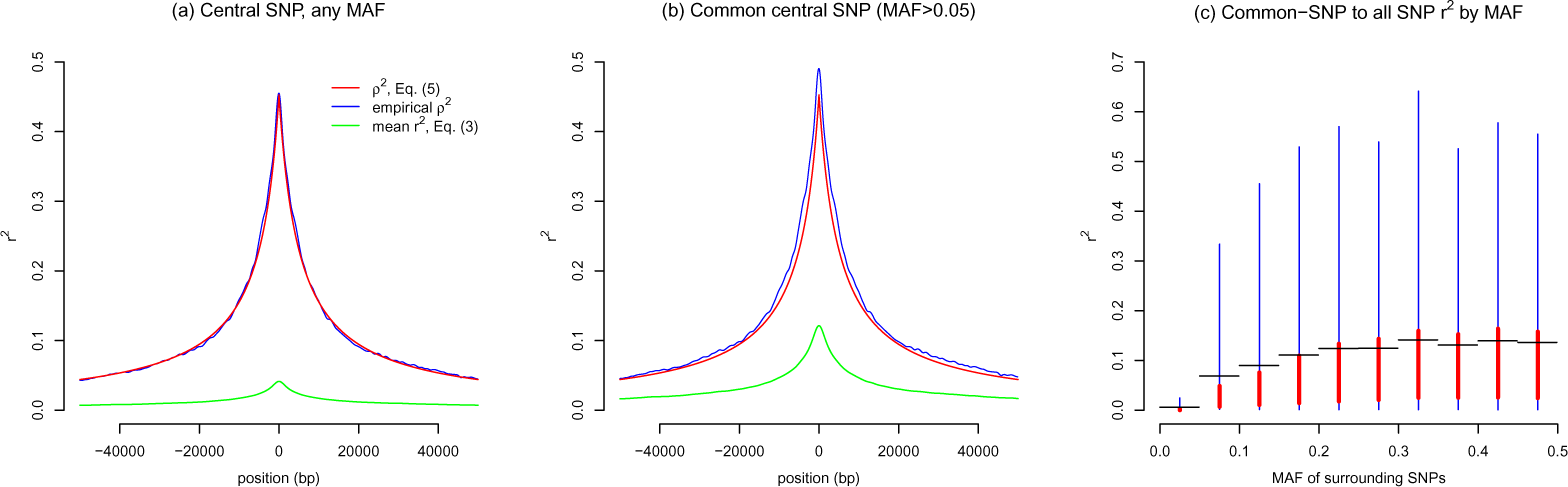
Mean *r*^2^ values between the hypermutable locus and SNPs with varying MAF. The mean *r*^2^ values are represented by the horizontal black line. The top (bottom) of the vertical red line represents the 75th (25th) percentile, and the top (bottom) of the thinner blue lines represents the 95th (5th) percentile. Results for simulations using a hypermutable mutation rate of 10^−3^ (a), 10^−4^ (b), and 10^−5^ (c) are shown.

In general, the SNP MAF only has a weak effect on the mean *r*^2^; the range of *r*^2^ values is similar for most SNP MAFs. However, for the lowest-MAF SNPs, the maximum possible *r*^2^ values are very small and the distribution of *r*^2^ shows that almost all low-MAF SNPs have very weak associations with the hypermutable locus. More importantly, Fig. 3 (b) and (c) show that common SNPs (MAF ¿ 0.1) can sometimes be in relatively high LD (*r*^2^ > 0.2) with hypermutable loci at the lower range of mutation rates (*μ* = 10^−4^ to 10^−5^).

### 1.3 SNP-SNP correlations

To put all of the above results in context, we examine how SNPs are correlated with each other. We find that, on average, SNPs have an extremely low mean *r*^2^ value with other SNPs (Fig. 4 (a)). The maximum mean *r*^2^ value, provided by SNPs in close proximity to the central SNP, is less than 0.05. Importantly, most SNPs have extremely low MAF (Fig. 1(d)), and the mean *r*^2^ value is strongly influenced by weak associations with rare SNPs (not shown). The correlation between common SNPs and rare SNPs is known to be weak [40], so the lack of a regional association between a single rare SNP and surrounding SNPs is expected. Furthermore, this scenario represents a breakdown of the approximation; the value of N(c+k) is too small for the approximation to be accurate. Therefore the predicted and emperical *ρ*^2^ of almost 0.45 for the SNPs that are in close proximity are clearly not a good approximation for the mean *r*^2^.

**Figure 4.**
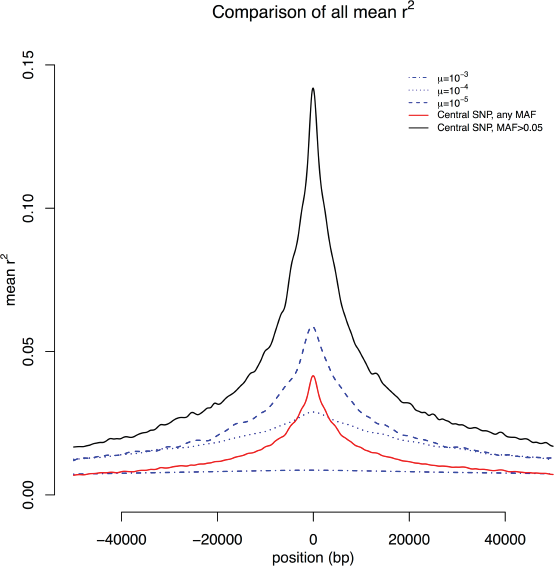
The *r*^2^ between a central SNP and surrounding SNPs. (a) Mean *r*^2^ values for SNP-SNP pairs, using a central SNP with any MAF (green). Also the analytical approximation for *r*^2^ (*ρ*^2^, red), and empirical *ρ*^2^ (blue). (b) Same as in (a) but for central SNPs with an MAF above 0.05, i.e. common central SNPs. (c) Distribution of *r*^2^ for comparisons between a central SNP with MAF above 0.05 and surrounding SNPs binned by their MAF. The mean *r*^2^ values are represented by the horizontal black line. The top (bottom) of the vertical red line represents the 75th (25th) percentiles, and the top (bottom) of the thinner blue lines represent the 95th (5th) percentiles.

Because hypermutable elements tend to have higher MAFs, perhaps a more appropriate comparison is to examine a central SNP only if its MAF is above 0.05. When these common central SNPs are examined for their correlations with surrounding SNPs with any MAF, the mean *r*^2^ values increase, but again the approximation (5) is not a good approximation for *E* (*r*^2^) because again N(c+k) is too small (Fig. 4(b)). To explore how the MAF of surrounding SNPs affects these values, we plot the *r*^2^ values for correlations between a central common SNP and surrounding SNPs with binned MAF (Fig. 4 (c)). Again the rare SNPs (MAF ¡ 0.05) show a very weak association, and common SNPs show a higher correlation. Intriguingly, common SNPs tag rare SNPs worse than they tag (the often common) hypermutable elements.

The correlations found using common central SNPs are similar to those found with hypermutable elements with a mutation rate of 10^−5^ (Fig. 2). However, the distribution of the *r*^2^ values for common central SNPs (Fig. 4 (c)) indicates that the upper 95th percentile of *r*^2^ values for common SNP associations are higher than those of any hypermutable element (Fig. 3 (c)). Therefore, large *r*^2^ values (e.g.*r*^2^ > 0.5) will be more frequent between common SNPs than between any hypermutable element and surrounding SNPs.

### 1.4 Relating hypermutable locus-SNP correlations with SNP-SNP correlations

To compare the mean *r*^2^ values for each scenario used, we plot all of the mean *r*^2^ values for all simulations together (Fig. 5). This plot demonstrates the relatively high mean *r*^2^ values for common SNPs (peaking just below 0.15), and a lower mean *r*^2^ values for loci with a mutation rate of 10^−5^. Additionally, loci with a mutation rate of 10^−4^ provide an interesting comparison to the analysis using all SNPs. In close proximity, the mean *r*^2^ measured on all SNPs is higher than that for loci with a mutation rate of 10^−4^, but the correlation decays with distance much more rapidly for the SNPs. At a distance of 4000 bp the mean *r*^2^ is nearly zero for all SNPs, but it remains above 0.1 at 4000 bp for hypermutable loci with mutation rates of 10^−4^ and 10^−5^.

**Figure 5.**
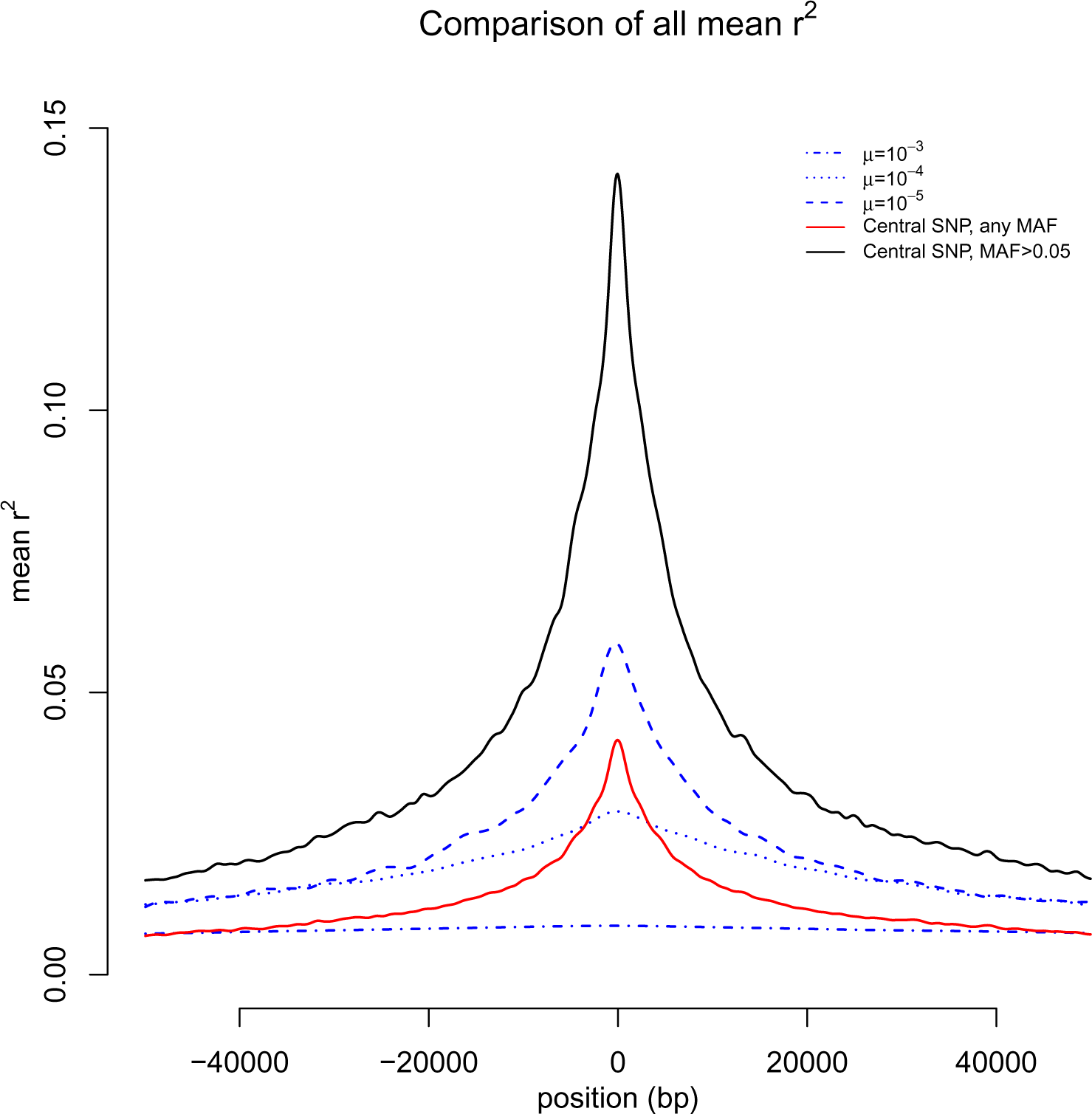
The mean *r*^2^ between surrounding SNPs, with any MAF, and a central variant. Various different central loci were examined: mutation rates of 10^−3^, 10^−4^, and 10^−5^, as well as a central SNP with any MAF and also a central common SNP (MAF¿0.05).

To further investigate these simulation results, we examine the locus with the largest *r*^2^ found in each simulation, 2000 simulations per scenario. The maximum *r*^2^ that occurs in an individual population is of interest because GWAS associations typically focus on SNPs with the lowest p-values. The scatter plot of the maximum per-simulation *r*^2^ for a central hypermutable locus (Fig. 6(a)) demonstrates that SNPs with the strongest associations are more centralized in the simulations using lower mutation rates than in those using higher mutation rates. There is almost no localization in the simulations with *μ* = 10^−3^ (Fig. 6 (c)). Furthermore, the maximum *r*^2^ values under the mutation rate of 10^−3^ are always small; the largest maximum *r*^2^ was only 0.202.

**Figure 6.**
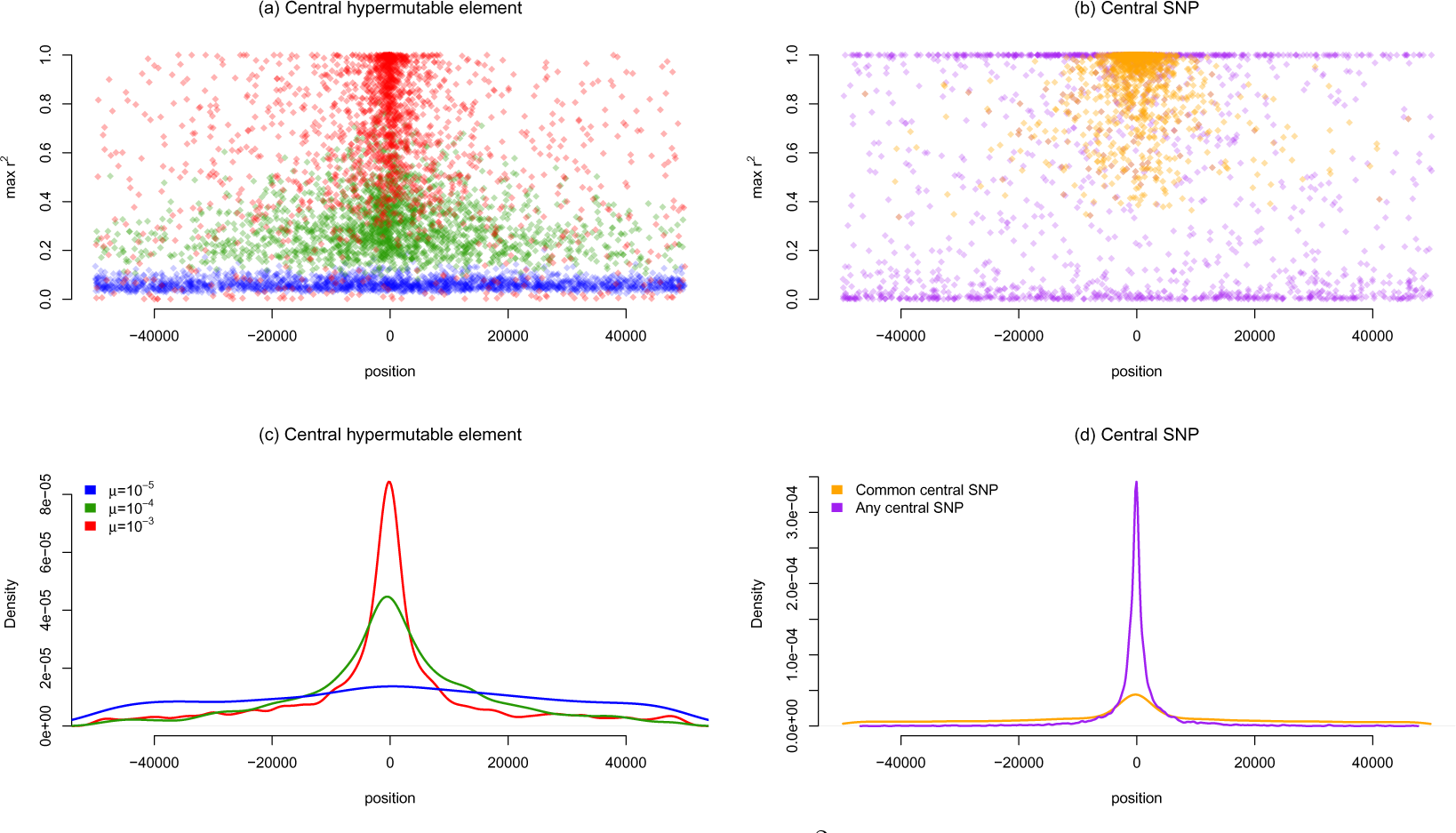
Characteristics of the maximum *r*^2^ between a central element and surrounding SNPs from each individual simulation, 2000 in total. (a) Scatterplot of the maximum *r*^2^ against the position relative to a central hypermutable element. Colors indicate mutation rate of the hypermutable element. (b) Scatterplot of the maximum *r*^2^ against the position relative to a central SNP (i.e., a central locus with normal mutation rate). Colors indicate MAF of the central SNP (common or unconstrained). (c) Density of the position of the locus with maximum *r*^2^, relative to a central hypermutable element. (d) Density of the position of the locus with maximum *r*^2^, relative to a central SNP.

When the central locus is a common SNP, the maximum *r*^2^ values are often near one (Fig. 6 (b)). When the central SNP is rare, the maximum *r*^2^ for the simulation is usually either very low or near one. Rare SNPs often have no association with surrounding loci, but occasionally a rare central SNP will be in perfect LD with another rare SNP, and this surrounding SNP in perfect LD is sometimes at a great distance. The maximum *r*^2^ for common central SNPs is often relatively large and localized to the central region (Fig. 6(d)).

## 2 Discussion

### 2.1 Comparing results from the approximation with simulations

The approximation made by [2], *E*(*r*^2^) ≈ 1/[4*N*(*c* + *k*)], provides a useful way to think about how mutation rates are related to linkage: the effects of mutation are similar to the effects of recombination, breaking linkage between loci. Although this approximation is only accurate when N(c+k) is large, one can nevertheless use it to build intuition about how mutation reduces correlations between loci. A forward-backward mutation rate of 10^−3^ acts like a genetic distance of 0.2 cM, about 200kb in humans (*k* ≈ 2*μ* = 0.002, corresponding to c = 0.002). Loci at a distance of 200kb are essentially unlinked. Therefore, even SNPs in close proximity to a hypermutable element with such a high mutation rate will be unlinked in genetic data. The best chance for a SNP to tag such a hypermutable element would be if the effective population size were small. This simple approximation makes it clear that SNPs do not tag variation caused by the most hypermutable loci in the human genome, except perhaps in highly inbred populations. Furthermore, the simulations demonstrate that the approximation of [2] over-estimates *E* (*r*^2^). When a site mutates rapidly, almost all of its correlation with surrounding loci is lost.

The approximation breaks down when N(c+k) is smaller than one [2], which is the case for most of the scenarios examined here. In these scenarios, the ratio of expectations in (4), *ρ*^2^ is a poor approximation for the expectation of the ratio given in (3). The only scenario in which *N* (*c* + *k*) is larger than one is when the mutation rate is 10^−3^ (Fig. 2 (c)). Oddly, this is also the only scenario in which empirical *ρ*^2^ does not appear to match the analytical approximation *ρ*^2^ of equation (5).

Therefore, although the approximation made by [2] can be helpful for understanding how mutation rates relate to recombination distance, simulations are required to estimate the mean *r*^2^ values for hypermutable elements with mutation rates larger than 10^−3^. For investigating these mutation rates, neither decreasing the population size nor increasing genetic distance would increase the accuracy or utility of the approximation. The diffusion approximation breaks down as population sizes decrease. Furthermore, our interest here is to understand how SNPs can tag nearby hypermutable elements, and examining SNPs that are a great distance to a hypermutable element provides limited utility because a tiny *r*^2^ is expected across large genetic distances. Thus the approximation *ρ*^2^ has many limitations when studying hypermutable elements.

The simulation results provide useful insight into how SNPs correlate with hypermutable elements. For most hypermutable elements, the mean *r*^2^ values with nearby SNPs are small, especially in comparison to common SNP-SNP associations (Fig. 5). However, for hypermutable elements with mutation rates of 10^−5^ not all of the correlation is lost. The mean *r*^2^ value for mutation rates of 10^−5^ is approximately half that of common SNP-SNP associations (Fig. 5). Furthermore, for a mutation rate of 10^−5^ the top 5th percentile of *r*^2^ values are all above 0.3 when the surrounding SNPs have an MAF above 0.2 (Fig. 3(c)). Stronger associations exist between common SNPs and other common SNPs (Fig. 4 (c)), but the scenario with mutation rates of 10^−5^ is somewhat comparable.

Rare SNPs are known to have a small *r*^2^ value with other SNPs [40], and rare SNPs are a potential explanation for missing heritability [24] and still-missing heritability [25, 26]. The simulations indicate that rare SNPs have a low mean *r*^2^ with other SNPs, comparable to hypermutable elements with mutation rates of 10^−4^ or smaller. However, the mean *r*^2^ diminishes across genetic distance faster for SNPs than for hypermutable loci (Fig. 5). This suggests that although hypermutable elements may behave similarly to rare SNPs, associations with hypermutable elements may show weaker localization. This delocalization spreads associations with hypermutable loci around the genome. Therefore, methods that use all SNPs together to measure overall genetic effects, such as GCTA [22], may be able to recover information about causal hypermutable loci.

### 2.2 Implications for GWAS

Hypermutable tandem repeat loci may be partially responsible for missing heritability [18, 27] and also still-missing heritability. The results presented here suggest that loci with high mutation rates are not well tagged by SNPs, and therefore much of the heritable variation caused by such loci will not have been captured in modern GWAS analyses. Scientists have just recently begun to estimate the mutation rates of hypermutable elements in the human genome [8–10, 15], and a database of known tandem repeat variants has recently been developed [4]. As more tandem repeat variants are cataloged, understanding how these variants can be tagged by SNPs will allow researchers to measure their relative contributions to phenotypes.

An important consideration when investigating a GWAS signal is the distance between the SNP with the lowest p-value and the variant(s) driving the association. The position of the lowest p-value SNP is often used to link a gene with a phenotype. Our results suggest that the top SNP associations are far less localized for hypermutable elements, with almost no localization for elements with a mutation rate of 10^−3^ (Fig. 6 (c)). Therefore, if a hypermutable element is causing a SNP association, the strongest SNP association may occur at a great distance from the causal element. Associations with hypermutable elements are also spread across a larger region (Fig. 5), providing an association signature that may be noticeably distinct from other types of associations.

Finally, because traits can be influenced by hypermutable elements and/or low frequency variants, SNP data alone cannot be used to exclude a gene or region of the genome as causal. If a gene is affected by hypermutable elements and/or rare variants, then SNPs will often fail to find an association. Regions or genes that contain potentially functional hypermutable elements require further genotyping of these elements before they can be totally excluded as potentially impacting a trait. Furthermore, many sequencing technologies have a limited ability to genotype some tandem repeat variants [27, 41–44], so our results apply to any data that is limited to SNPs. Recent advances in sequencing technology [42, 43, 45] and tandem repeat genotyping [6, 44, 46, 47] provide hope that some hypermutable elements will be included in future studies of genetic heritability and genetic disease. Nevertheless, some of the missing heritability caused by hypermutable elements may remain missing, at least for the near future.

### 2.3 Limitations and potential extensions

This study uses only two possible states at each locus, and the forward and backward mutations are equal. This simplifies both the analytical approach as well as the simulations, and can be used as a simple model of tandem repeat evolution. Tandem repeats often have more than two states, but diseases caused by tandem repeats are often caused by expansion [14, 18]. Therefore, tandem repeats can sometimes fit into a two-allele model as was done here (short versus long). However, transitions between a short allele and a long allele depend on the repeat length [8], and thus forward and backward mutation rates are not necessarily equivalent. A step-wise mutation model, allowing multiple allele sizes at the hypermutable locus and binning them as short or long, may provide a more accurate model of tandem repeat diseases. These more complicated models are likely to return similar results because empirical data indicate that small *r*^2^ values are found between SNPs and tandem repeat loci, whether they are bi-allellic or multi-allelic [4].

The use of a stable population with an effective size of 10,000 without population history may further limit the direct application of these results. The results from smaller population sizes might drastically change because the diffusion approximation does not work well for small effective population sizes. In addition, complicated population histories may change these results in unexpected ways, especially because tandem repeats and SNPs provide different information about population histories [48]. Future simulations could address these possibilities.

Equation (6) suggests that increasing the population size will result in an approximately harmonic decrease in the mean *r*^2^. Therefore, one can expect the mean *r*^2^ from an effective population size of 20,000 to be approximately half of the mean *r*^2^ found here with an effective population size of 10,000. Extrapolating the results presented here to smaller population sizes would not be as straightforward. Due to the aforementioned effect that small populations have on the accuracy of the diffusion approximation, estimating how these result would change if one used a smaller population size is not as simple as applying a linear transformation.

### 2.4 Summary and conclusion

As shown by [2], mutation and recombination act in a similar fashion to break up linkage between loci. The magnitude of the mutation rate can be approximately equated to recombination distance in centimorgans. However, this approximation only holds when the mutation rates are high and/or population sizes are large. With lower mutation rates the approximation breaks down and simulations must be used to estimate the expected linkage between loci.

The simulations reported here suggest that the variation caused by some hypermutable elements can be captured using SNPs. At mutation rates of 10^−5^ or smaller the associations between hypermutable loci and SNPs is comparable to, although lower than, common SNP - common SNP associations. On the other hand, the correlations between SNPs and loci with mutation rates of 10^−4^ and 10^−3^ are relatively low, and therefore variation caused by loci with these mutation rates are likely to show only weak association with SNPs of any MAF.

Heritable variation can be caused by genetic loci with a range of mutation rates [1]. Hypermutable loci can remain highly polymorphic in a population, and they may be important causes of human disease and heritability of complex traits. Common SNP variants are currently inexpensive and widely used to search for genes that contribute to heritable variation. Unfortunately, many hypermutable loci will have poor linkage with SNPs, and therefore these loci will be unlikely to be uncovered using SNP GWAS methods. Direct genotyping will be necessary to uncover the effects that many hypermutable loci have on genetic variation. We hope that this work will help researchers investigating the sources of human diseases and heritable traits.

